## Supplementary figures and images for "Spatiotemporal patterns of neocortical activity around hippocampal sharp-wave ripples"

### Supplemental Data 1

*i.*

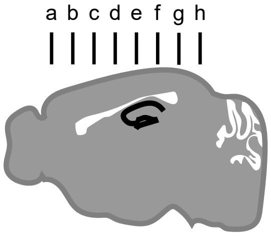


*ii.*

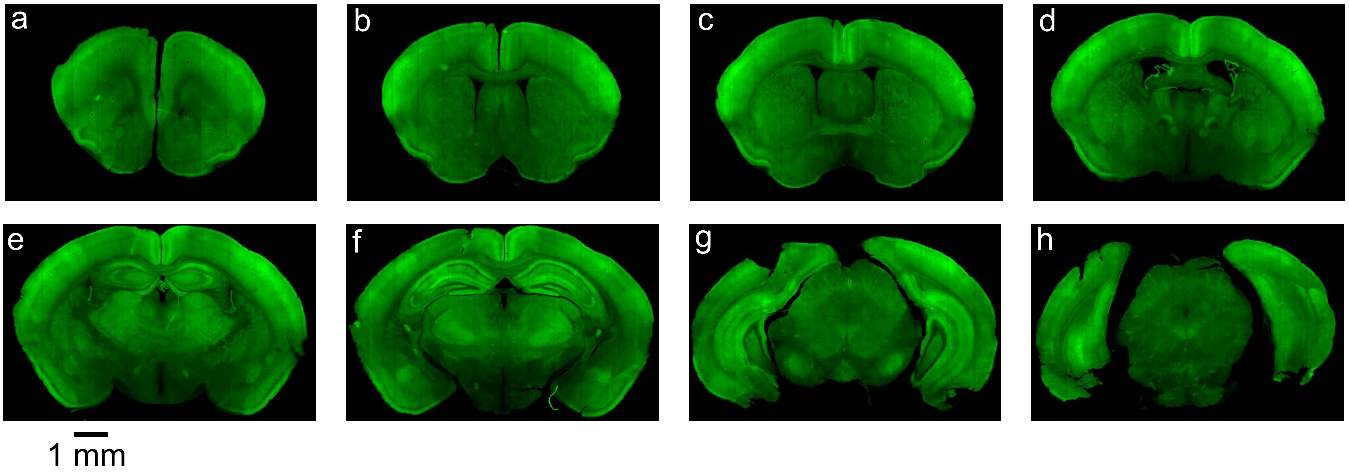


*iii.*

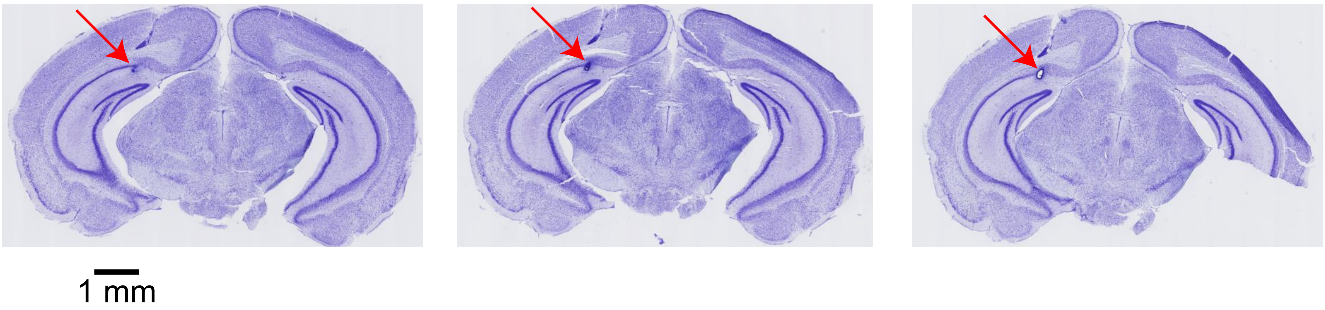


*iv.*

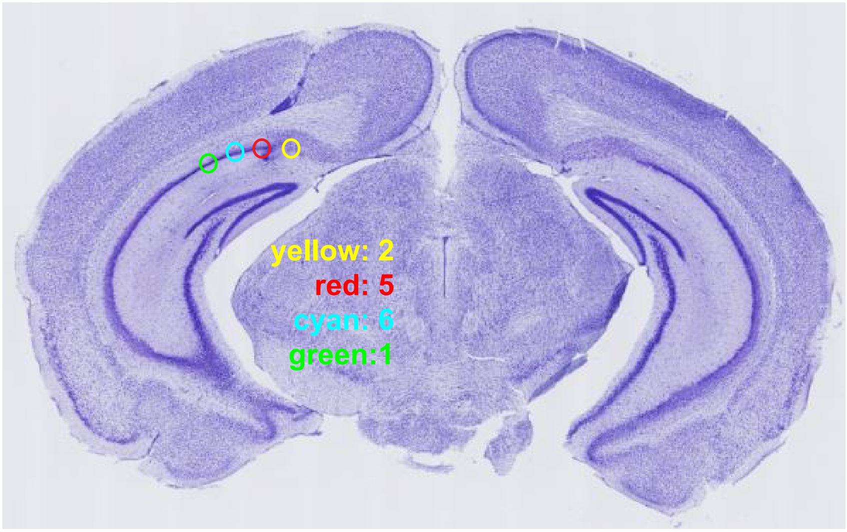

### Supplemental Data 2

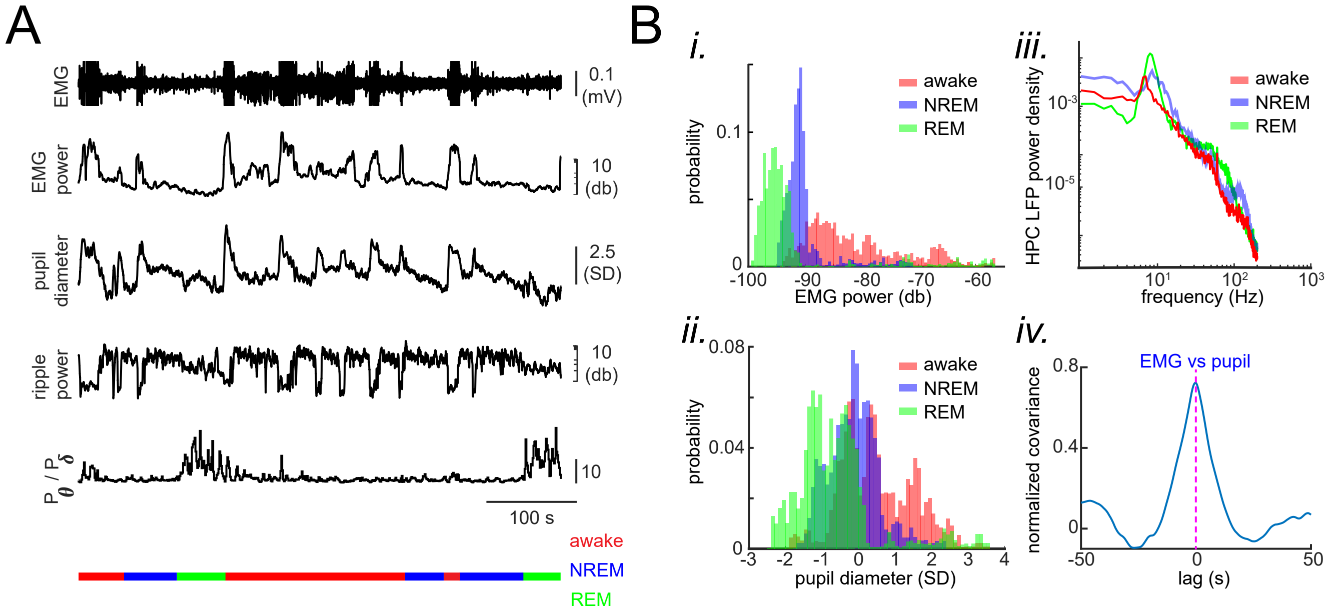

### Supplemental Data 3

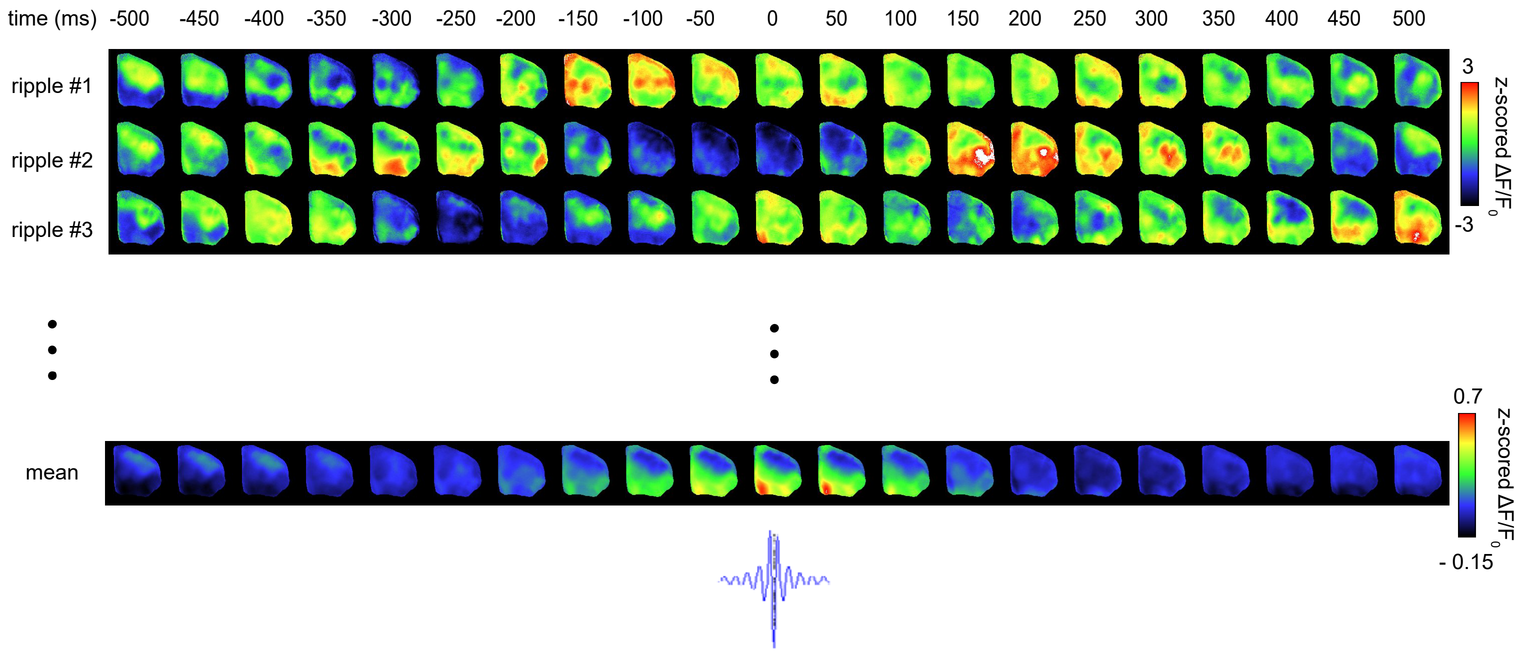

### Supplemental Data 4

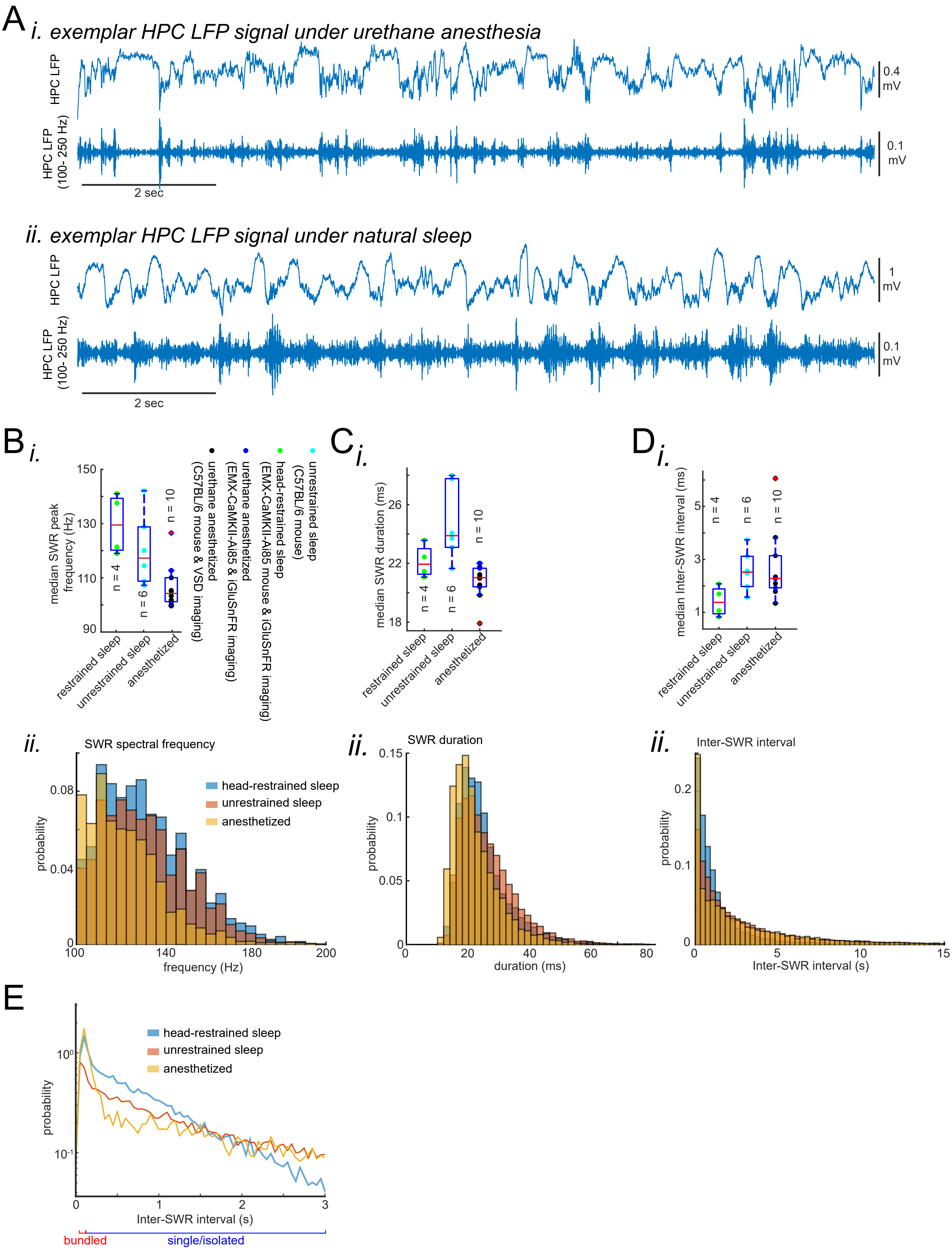

### Supplemental Data 5

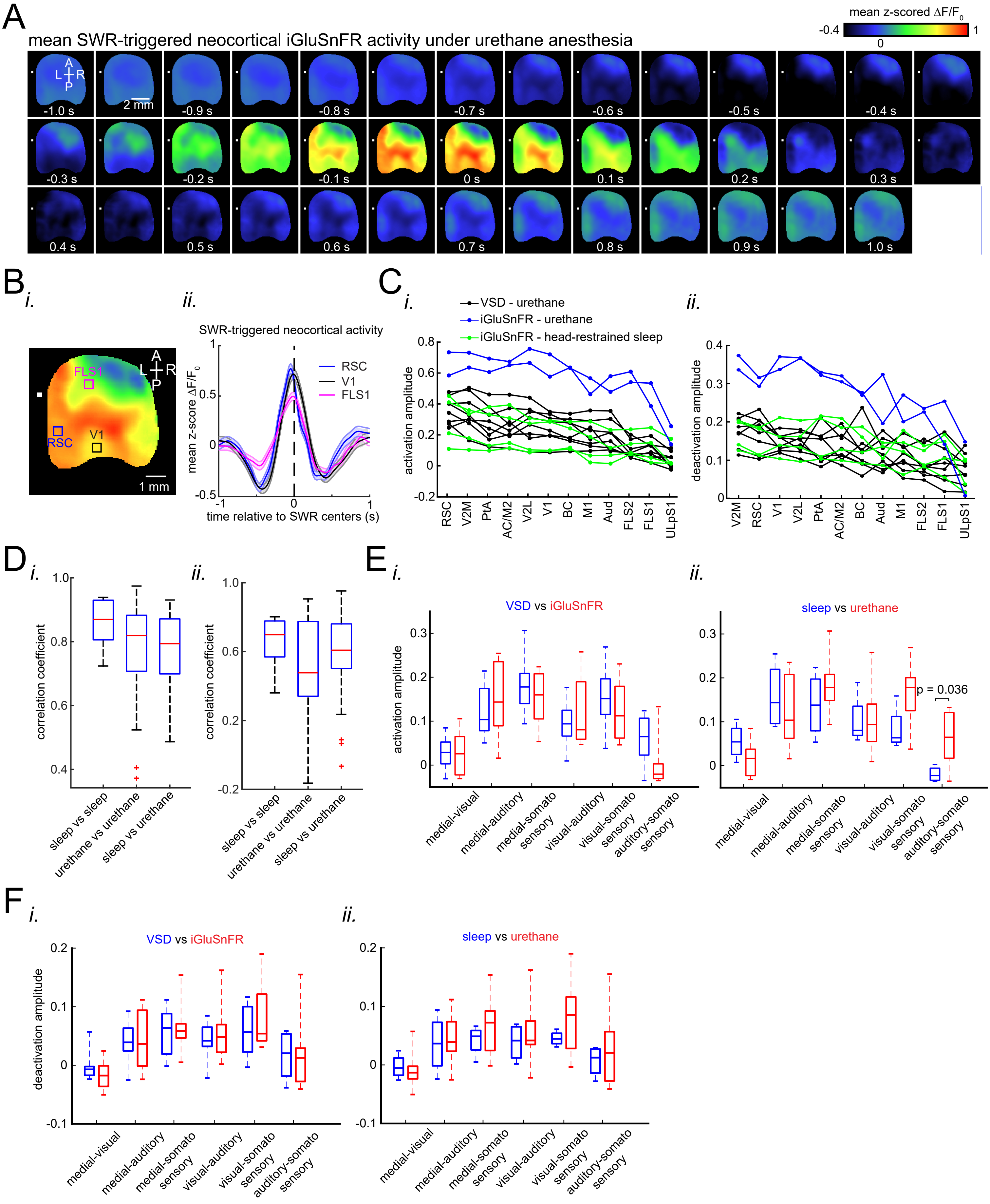

### Supplemental Data 6

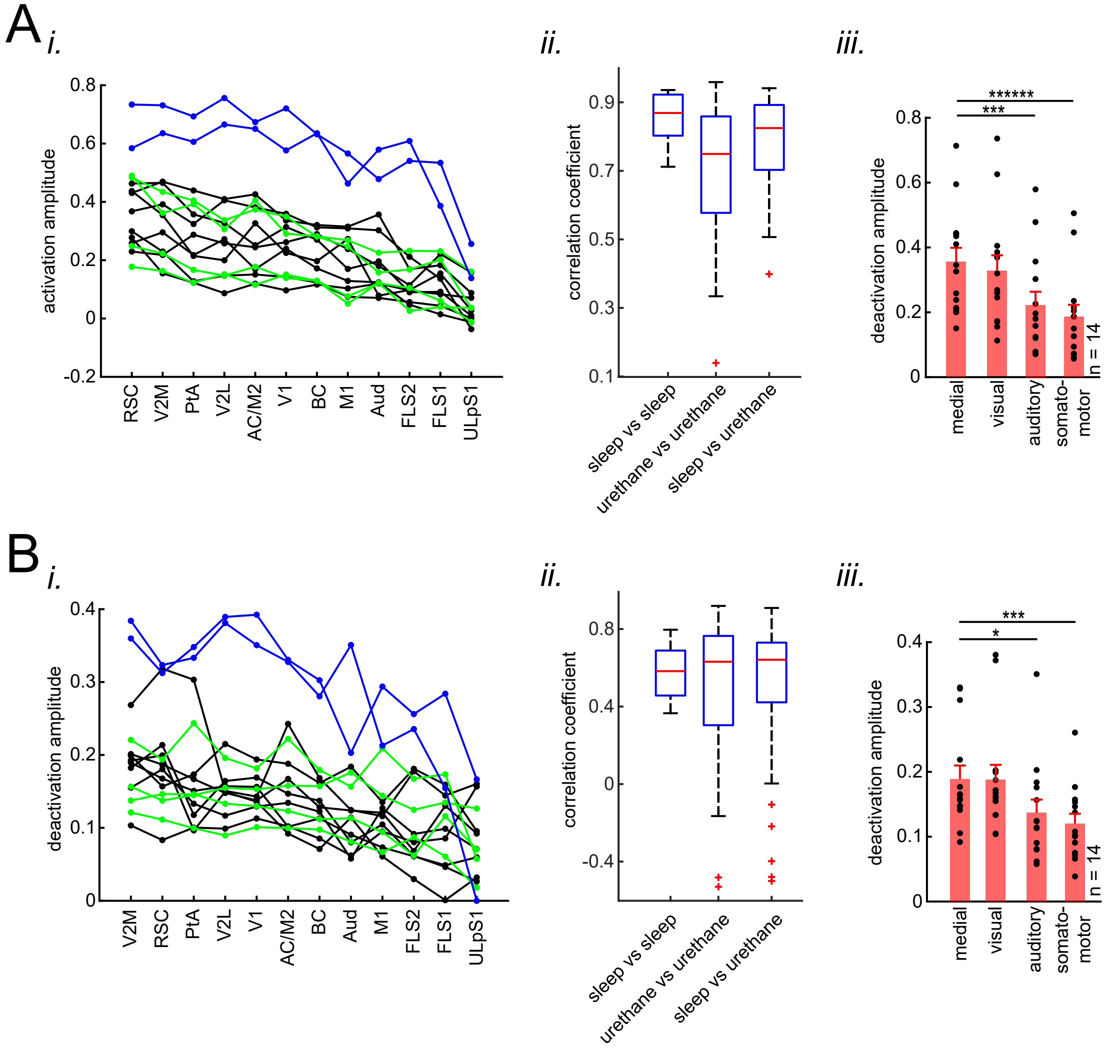

### Supplemental Data 7

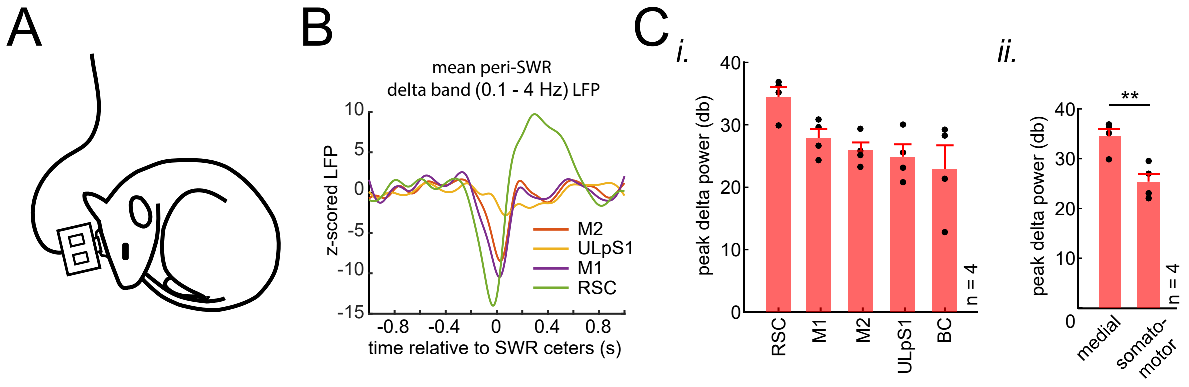

### Supplemental Data 8

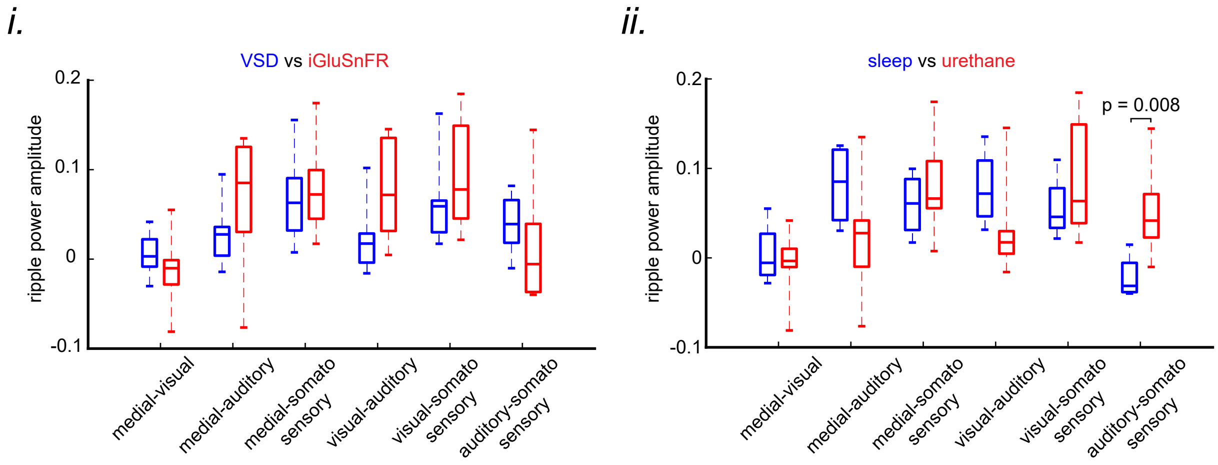

### Supplemental Data 9

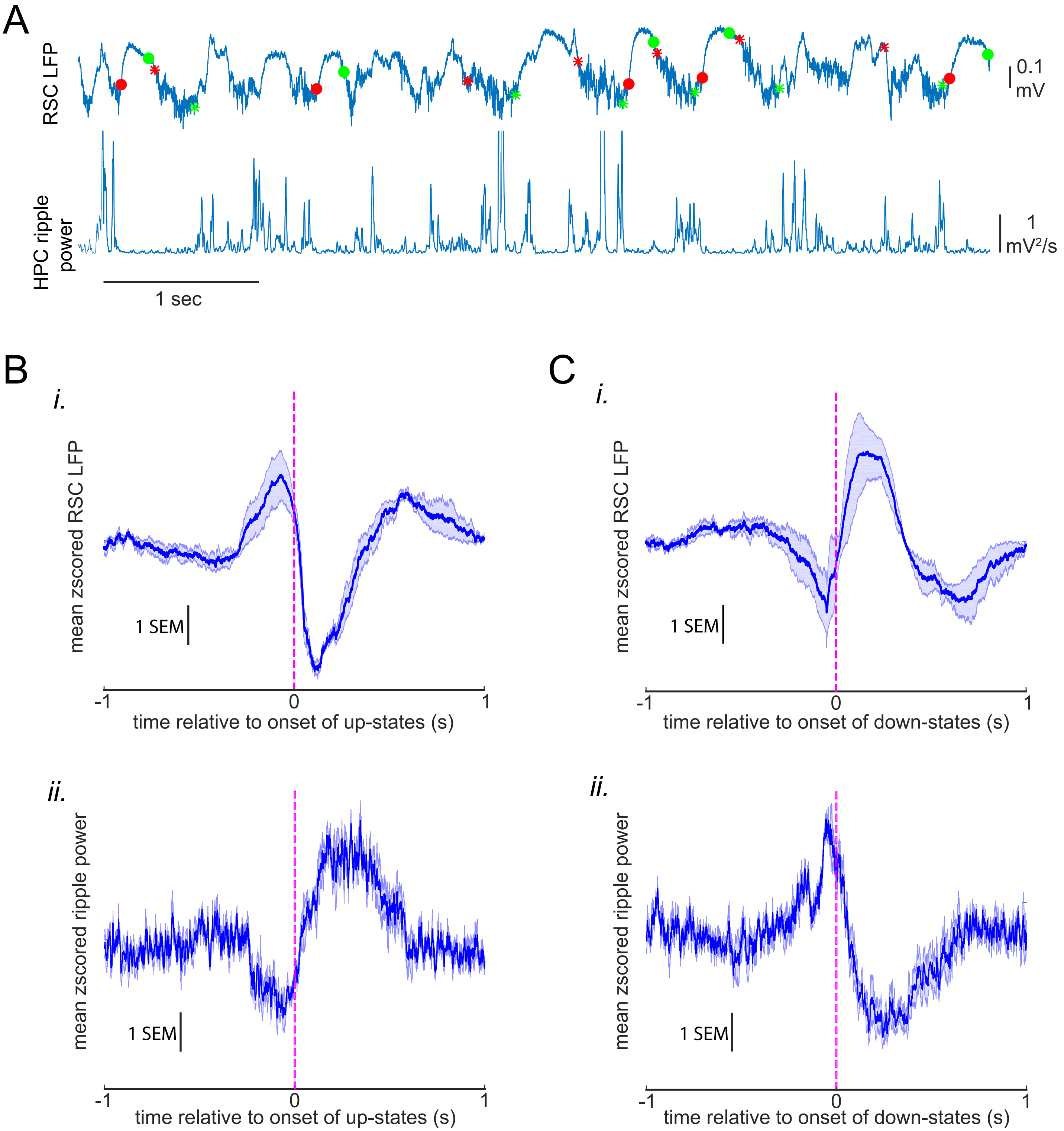

### Supplemental Data 10

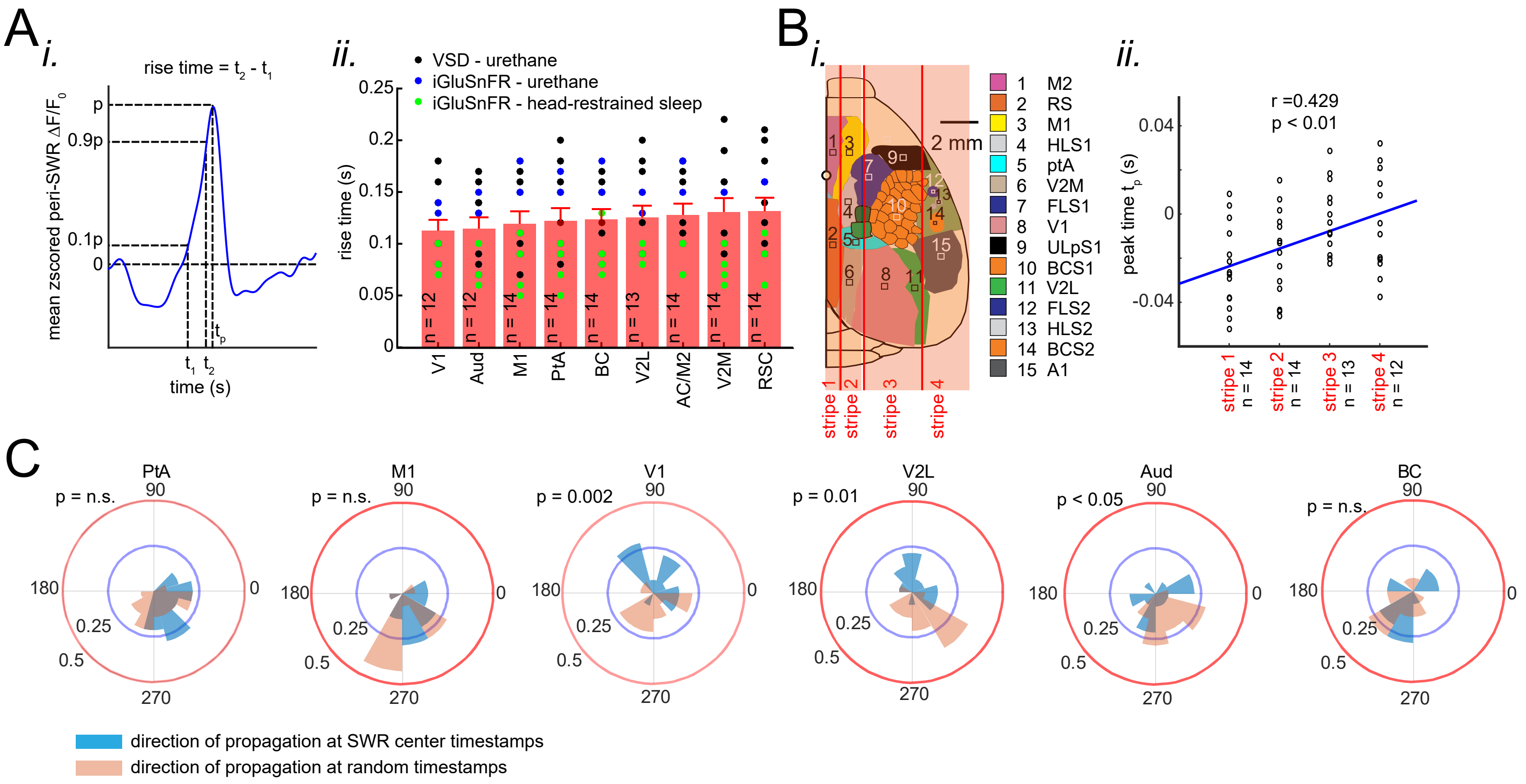

### Supplemental Data 11

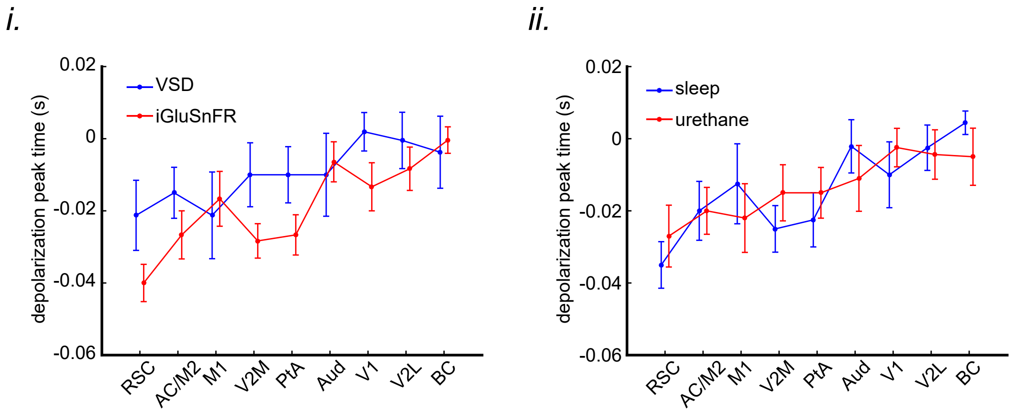

### Supplemental Data 12

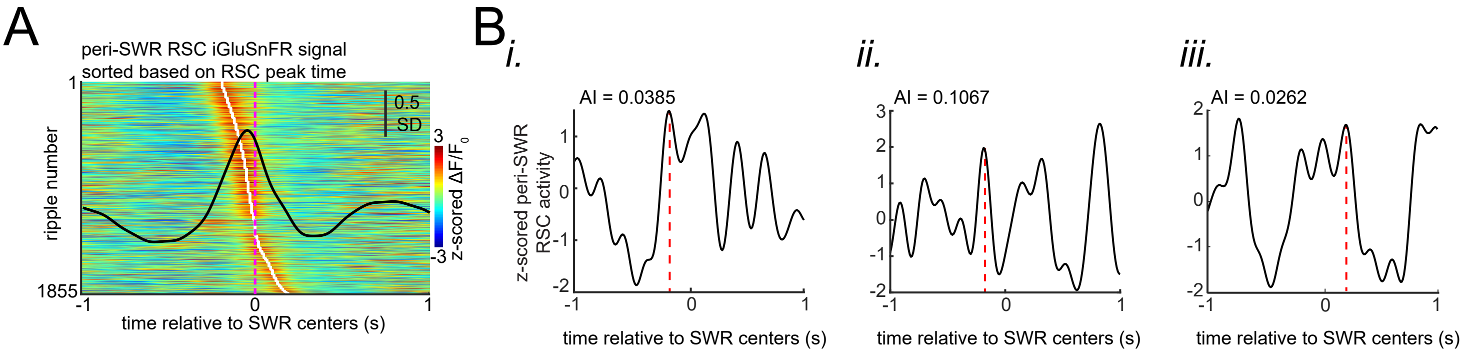

### Supplemental Data 13

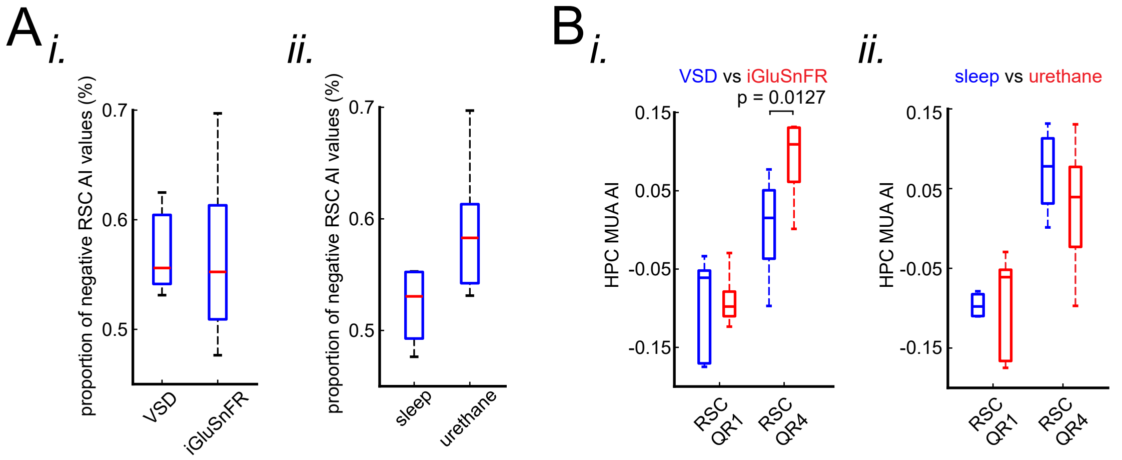

### Supplemental Data 14

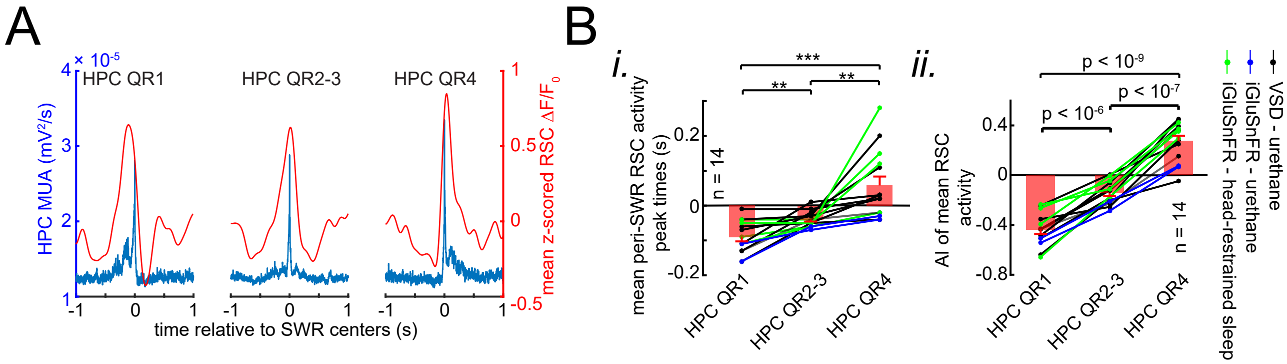

### Supplemental Data 15

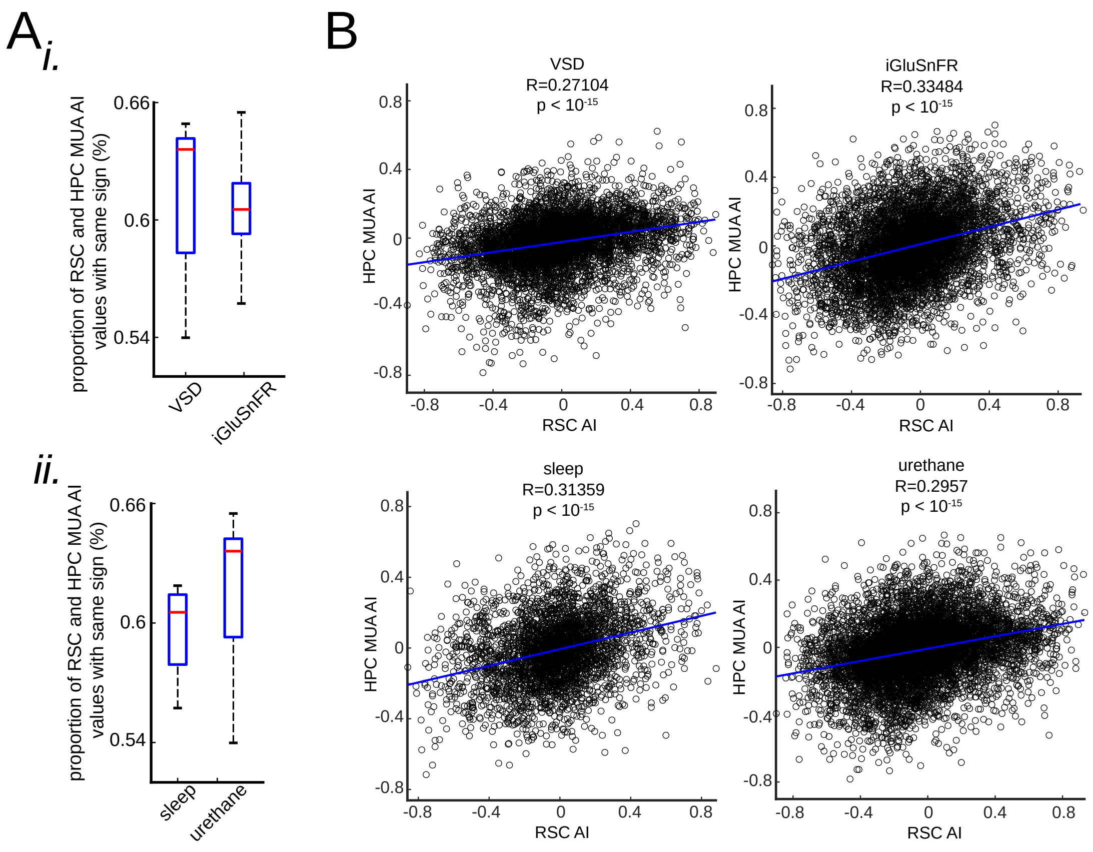

### Supplemental Data 16

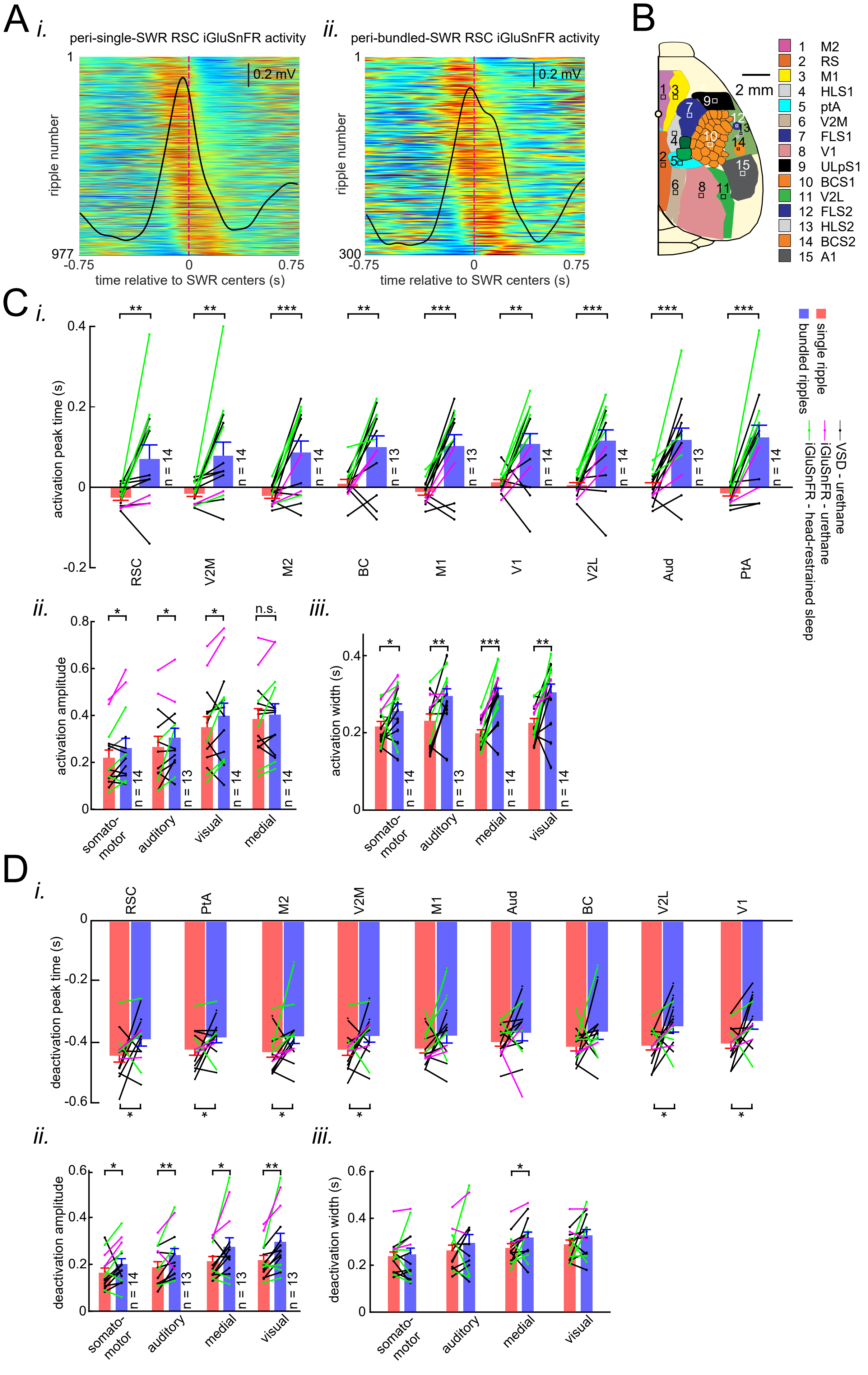

### Supplemental Data 17

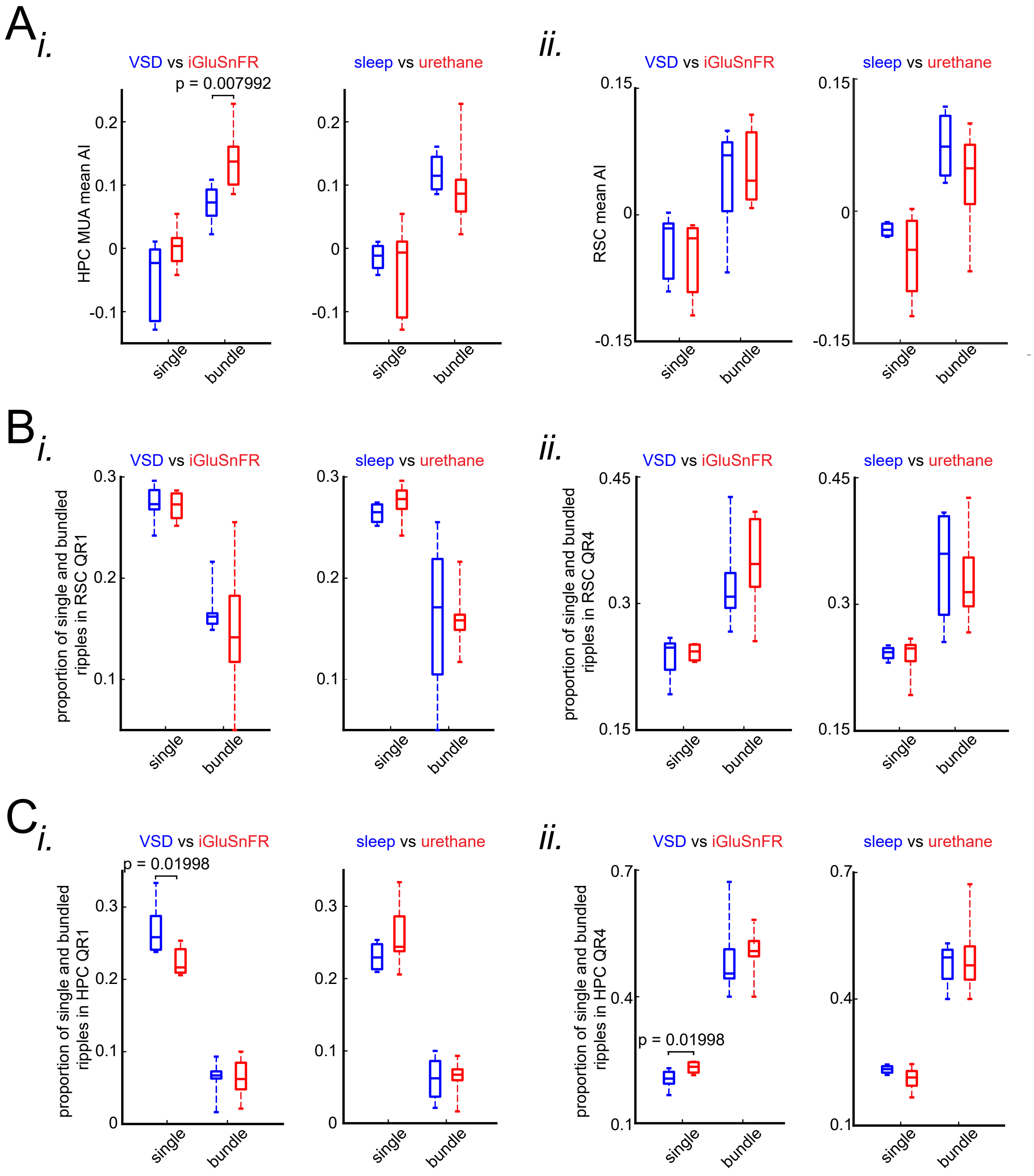
